# Spatial dynamics within and between brain functional domains: A hierarchical approach to study time-varying brain function

**DOI:** 10.1101/391094

**Authors:** A. Iraji, Z. Fu, E. Damaraju, T.P. DeRamus, N. Lewis, J.R. Bustillo, R.K. Lenroot, A. Belger, J.M. Ford, S. McEwen, D.H. Mathalon, B.A. Mueller, G.D. Pearlson, S.G. Potkin, A. Preda, J.A. Turner, J.G. Vaidya, T.G.M. van Erp, V.D. Calhoun

## Abstract

The analysis of time-varying activity and connectivity patterns (i.e., the chronnectome) using resting-state magnetic resonance imaging has become an important part of ongoing neuroscience discussions. The majority of previous work has focused on variations of temporal coupling among fixed spatial nodes or transition of the dominant activity/connectivity pattern over time. Here, we introduce an approach to capture spatial dynamics within functional domains (FD), as well as temporal dynamics within and between FD. The approach models the brain as a hierarchical functional architecture with different levels of granularity, where lower levels have higher functional homogeneity and less dynamic behavior and higher levels have less homogeneity and more dynamic behavior. First, a high-order spatial independent component analysis is used to approximate functional units. A functional unit is a pattern of regions with very similar functional activity over time. Next, functional units are used to construct FDs. Finally, functional modules (FMs) are calculated from FDs, providing an overall view of brain dynamics. Results highlight the spatial fluidity within FDs, including a broad spectrum of changes in regional associations from strong coupling to complete decoupling. Moreover, FMs capture the dynamic interplay between FDs. Patients with schizophrenia show transient reductions in functional activity and state connectivity across several FDs, particularly the subcortical domain. Activity and connectivity differences convey unique information in many cases (e.g. the default mode) highlighting their complementarity information. The proposed hierarchical model to capture FD spatiotemporal variation provides new insight into the macroscale chronnectome and identifies changes hidden from existing approaches.

## 1. Introduction

Neuronal populations interact with each other at different spatial scales (from micro to macro). At the macroscale, studying functional interactions using functional magnetic resonance imaging (fMRI) has significantly enhanced our knowledge of brain functional systems. Examining functional connectivity across the brain using univariate and multivariate analyses has revealed replicable, large-scale brain networks, also known as functional domains (FDs). Alterations of FDs may be significantly associated with different physiological and psychological conditions (Arbabshirani et al., 2017; Garrity et al., 2007; Greicius, 2008; Menon, 2011; Seeley et al., 2009; Sorg et al., 2007). Each FD is comprised of a set of spatially distinct and temporally covarying functional units (subnetworks), which putatively orchestrate various brain functions (van den Heuvel and Hulshoff Pol, 2010). A functional unit can be defined as a pattern of regions with very similar functional activity over time given the associated imaging modality. Hierarchical models of brain function posit that the brain has different levels of functional granularity, where lower levels are associated with reduced complexity. In other words, lower levels of the hierarchy display less functional dynamic behavior and higher functional homogeneity (Blumensath et al., 2013; Felleman and Van Essen, 1991; Meunier et al., 2010; Zhou et al., 2006).

At the same time, given the dynamic nature of the brain, recent studies have focused on capturing the time-varying information of the blood oxygenation-level dependent (BOLD) signal (Calhoun et al., 2014; Hutchison et al., 2013; Preti et al., 2017). Several strategies have been proposed to study time-varying information of BOLD signal, but most can be divided into one of two major categories. The first identifies reoccurring temporal coupling among fixed spatial nodes/networks (Allen et al., 2014; Barttfeld et al., 2015; Chen et al., 2016; Damaraju et al., 2014; Hutchison et al., 2013; Leonardi et al., 2013; Sakoglu et al., 2010; Shine et al., 2016; Yaesoubi et al., 2018). The most common approach for this category is the sliding-window technique (Allen et al., 2014; Sakoglu et al., 2010). The second category extracts the moment-to-moment dominant spatial co-activation or connectivity pattern without capturing the spatiotemporal variations within and between functional organizations (Karahanoglu and Van De Ville, 2015; Liu et al., 2013; Liu and Duyn, 2013; Preti and Van De Ville, 2017; Tagliazucchi et al., 2012; Trapp et al., 2018; Vidaurre et al., 2017). The co-activation patterns (CAPs) approach and its derivatives are used most frequently within this category (Karahanoglu and Van De Ville, 2015; Liu et al., 2013).

However, these approaches do not capture the ongoing spatial variations of brain functional organization, such as FDs, over time. In early work, Kiviniemi et al. used sliding-window ICA and observed spatial variations in the default mode network (Kiviniemi et al., 2011). Different spatial patterns were also observed for CAPs of the posterior cingulate cortex and the intraparietal sulcus over time (Liu and Duyn, 2013). Ma et al., shows fluctuations in spatial couplings by measuring residual mutual information between spatial components derived from independent vector analysis (Ma et al., 2014). These findings justify the need for an approach to measure variations in spatial patterns of brain functional organization over time. Additionally, given that the brain reorganizes its activity at different interacting spatial and temporal scales, investigating spatial dynamics (spatiotemporal variations) within and between different spatial scales provides a broader perspective of how the brain naturally functions. Here, we propose a novel, data-driven approach to capture and characterize both the spatiotemporal variations of FDs, and the dynamic interactions between them. The approach utilizes the concept of the functional hierarchy and encapsulates the spatiotemporal variations of each FD from its associated functional units. We suggest high-order intrinsic connectivity networks (hICNs) obtained from a high-order spatial independent component analysis (ICA) are good approximations of functional units of macroscale brain communication.

Using hICNs, we construct the elements of the higher hierarchical level (i.e., FDs) and study their spatial dynamics. Our findings highlight that FDs spatially evolve over time, (i.e., spatially vary over time). We characterize highly reproducible and distinct activity patterns called spatial domain states, within each FD. At various times, the interactions within and between FDs involve different spatial regions of the brain. Furthermore, evaluating the associations between FDs (i.e., functional state connectivity) identified distinct coupling patterns, called functional modules (FMs). FMs represent the transient patterns of temporal coupling between FDs and provide information of global brain temporal dynamics. One key advantage of the approach is its ability to successfully capture spatiotemporal changes of FDs, without applying constraints on their spatial and/or temporal couplings. The approach does not require a sliding-window technique, so it can capture the maximum temporal frequency variations in the temporal profile (Yaesoubi et al., 2018). Furthermore, it allows the detection of fluctuations in the spatial coupling of FDs up to the maximum spatial resolution existing in the data.

We further evaluate the clinical utility of our approach by studying alterations in spatial dynamics within patients with schizophrenia (SZ) relative to healthy controls. Schizophrenia is a functionally heterogeneous disorder which can include delusions, hallucinations, disorganized speech, disorganized or catatonic behavior, and negative symptoms (e.g. apathy, blunted affect) (Association, 2013). It has been suggested that schizophrenia is related to a reduced capacity to integrate information across different regions (Kahn et al., 2015; Stephan et al., 2006), which can lead to reduced functional connectivity (Damaraju et al., 2014; Kahn et al., 2015). However, previous work does not provide much information regarding how this reduced integration manifests. The application of our hierarchical approach to study the spatial dynamics of FDs could potentially identify underlying mechanisms that define how patients with SZ integrate information. Furthermore, the approach has the unique ability to detect nuanced transient alterations in the spatial patterns of FDs. While previous functional connectivity analyses report hypoconnectivity among patients with SZ, our approach is in line with this trend, but also detects transient reductions in the functional activity within specific FDs. Importantly, alterations in functional activity can occur in the absence of changes in functional connectivity and vice versa suggesting the complementarity of these two approaches. Furthermore, functional state connectivity, measured for the first time, displayed similar but also distinct differences between healthy controls and patients with SZ compared to previous functional connectivity analyses including decreased functional state connectivity between subcortical and somatosensory and somatomotor domains within the functional modules.

In summary, we introduce an innovative framework to shed new light on time-varying spatial characteristics of brain function. The approach provides the backbone for examining spatiotemporal variations of brain functional organizations and the hierarchy of spatial dynamics which can improve our understanding of brain function. Results also show the potential of further leveraging this time-varying behavior for characterizing mechanisms for clinical features in patient groups.

## 2. Materials and Methods

### 2.1. Glossary and Outline of the Approach

There have been many terms and jargon used to define the functional architectures of the brain, for example “network”, “circuit”, “module”, “domain”, and “system” have all been used to define the same functional structure. At the same time, each term can also refer to different functional structures across studies. For instance, the term “network” has referred to a collection of elements but at different levels from set of anatomically separated regions to a cluster of functionally homogeneous voxels to cell-specific regulatory pathways inside of neurons (Erhardt et al., 2011a; Petersen and Sporns, 2015). As described in (Erhardt et al., 2011a), the way to avoid confusion is to ensure that all terms are clearly defined, thus here we provide a glossary of key terms used throughout this paper (Figure 1).

**Figure 1.**
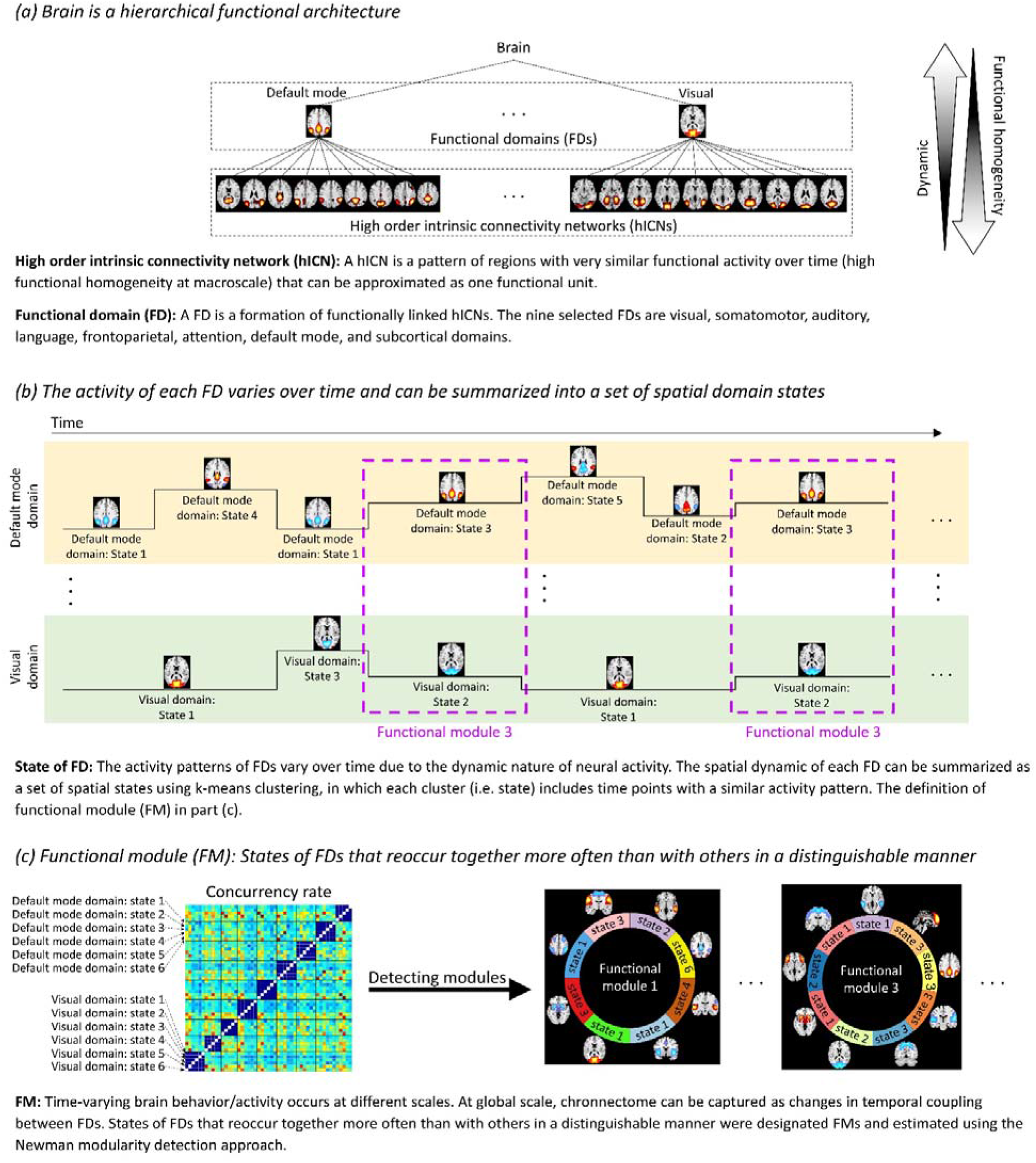
Hierarchical functional architectures of the brain and notations used in this work.

#### 1) High-order intrinsic connectivity network (hICN)

An hICN is comprised of a set of voxels (pattern of regions) with very similar functional activity over time (high functional homogeneity) that can be approximated as one functional unit. hICNs were obtained by applying high-order spatial ICA. The use of high-order ICA to generate hICNs instead of predefined anatomical locations allows us to detect functionally homogeneous regions from data itself (Calhoun and de Lacy, 2017).

#### 2) Functional domain (FD)

An FD is a formation of functionally linked hICNs. Focusing on cortical and subcortical regions, we define nine FDs based on a priori knowledge and results of low-order static spatial ICA. hICNs were categorized into FDs through a semi-automatic process. FDs, at a given point in time, were reconstructed from the associated hICNs and their time courses at that time point.

#### 3) State of FD

The spatial patterns of FDs vary over time due to the dynamic nature of neural activity. Spatial dynamics of FDs can be summarized as a set of spatial domain states using a clustering approach, in which each cluster (i.e. state) includes time points with a similar activity pattern.

#### 4) Functional module (FM)

The chronnectome occurs at different scales. At a global scale, the chronnectome can be captured by evaluating variations in temporal coupling of FDs. States of FDs that reoccur together more often than with others in a distinguishable manner are designated FMs, which are estimated using the Newman modularity detection approach.

### 2.2. Data Acquisition and Preprocessing

Data collection was performed at 7 imaging sites across the United States and passed data quality control. All participants were at least 18 years old and written informed consent was given prior to enrollment. Data was collected from 160 healthy controls including 46 females and 114 males (average age: 36.71 ± 10.92; range: 19-60 years) and 149 age- and gender-matched patients with SZ including 36 females and 113 males (average age: 37.95 ± 11.47; range: 18-60 years). Further details can be found in our earlier work (Damaraju et al., 2014).

MRI data were collected using a 3-Tesla Siemens Tim Trio scanner for six of the seven sites and on 3-Tesla General Electric Discovery MR750 scanner for the seventh site. Resting state fMRI data was collected using a standard gradient echo EPI sequence with following imaging parameters: pixel spacing size = 3.4375 × 3.4375 mm, FOV of 220 × 220 mm, matrix size = 64 × 64, slice thickness = 4 mm, slice gap = 1 mm, TR/TE = 2000/30 ms, flip angle = 77°, number of excitations (NEX) = 1, and acquisition time ≈ 5.4. During the resting state fMRI scans, participants were instructed to keep their eyes closed.

Data were preprocessed using a combination of SPM (http://www.fil.ion.ucl.ac.uk/spm/) and AFNI (https://afni.nimh.nih.gov) software packages including brain extraction, motion correction using the INRIAlign, slice-timing correction using the middle slice as the reference time frame, and despiking using AFNI’s 3dDespike. The data of each subject was subsequently registered to a Montreal Neurological Institute (MNI) template and resampled to 3 mm^3^ isotropic voxels, and spatial smoothed using a Gaussian kernel with a 6 mm full width at half-maximum (FWHM = 6 mm). Finally, voxel time courses were z-scored (variance normalized), as z-scoring has displayed improved parcellation of functional organizations structures (hICNs) compared to other scaling methods for independent component analysis.

### 2.3. High-order Intrinsic Connectivity Networks (hICNs) Extraction

ICA analysis was applied to obtain hICNs. Group ICA was performed using the GIFT software package from MIALAB (http://mialab.mrn.org/software/gift/) (Calhoun and Adali, 2012; Calhoun et al., 2001). First, data dimensionality reduction was performed using subject-specific spatial principal components analysis (PCA) followed by group-level spatial PCA (Erhardt et al., 2011b). The 200 principal components that explained the maximum variance were selected as the input for a high-order group-level spatial ICA to calculate 200 group independent components (Figure 2.A). High-order ICA allows us to segment the brain into a set of spatial patterns with very similar functional activity (high functional homogeneity) at the macroscale called hICNs (Allen et al., 2011; Kiviniemi et al., 2009). Infomax was chosen as the ICA algorithm because it has been widely used and compares favorability with other algorithms (Correa et al., 2007b; Correa et al., 2005). Infomax ICA was repeated 100 times using ICASSO framework (Himberg et al., 2004), and the ‘best run’ was selected to obtain a stable and reliable estimation (Calhoun and Adali, 2002; Calhoun et al., 2009; Correa et al., 2007a; Du et al., 2014; Ma et al., 2011). Sixty-five cortical and subcortical hICNs were selected and categorized into nine FDs based on their anatomical and common functional properties, and their relationships (spatiotemporal similarity) with independent components obtained from low-order spatial ICA (Figure 2.B). The selected hICNs should have peak activations in the gray matter and their time-courses be dominated by low-frequency fluctuations evaluated using dynamic range and the ratio of low frequency to high frequency power (Allen et al., 2011).

**Figure 2.**
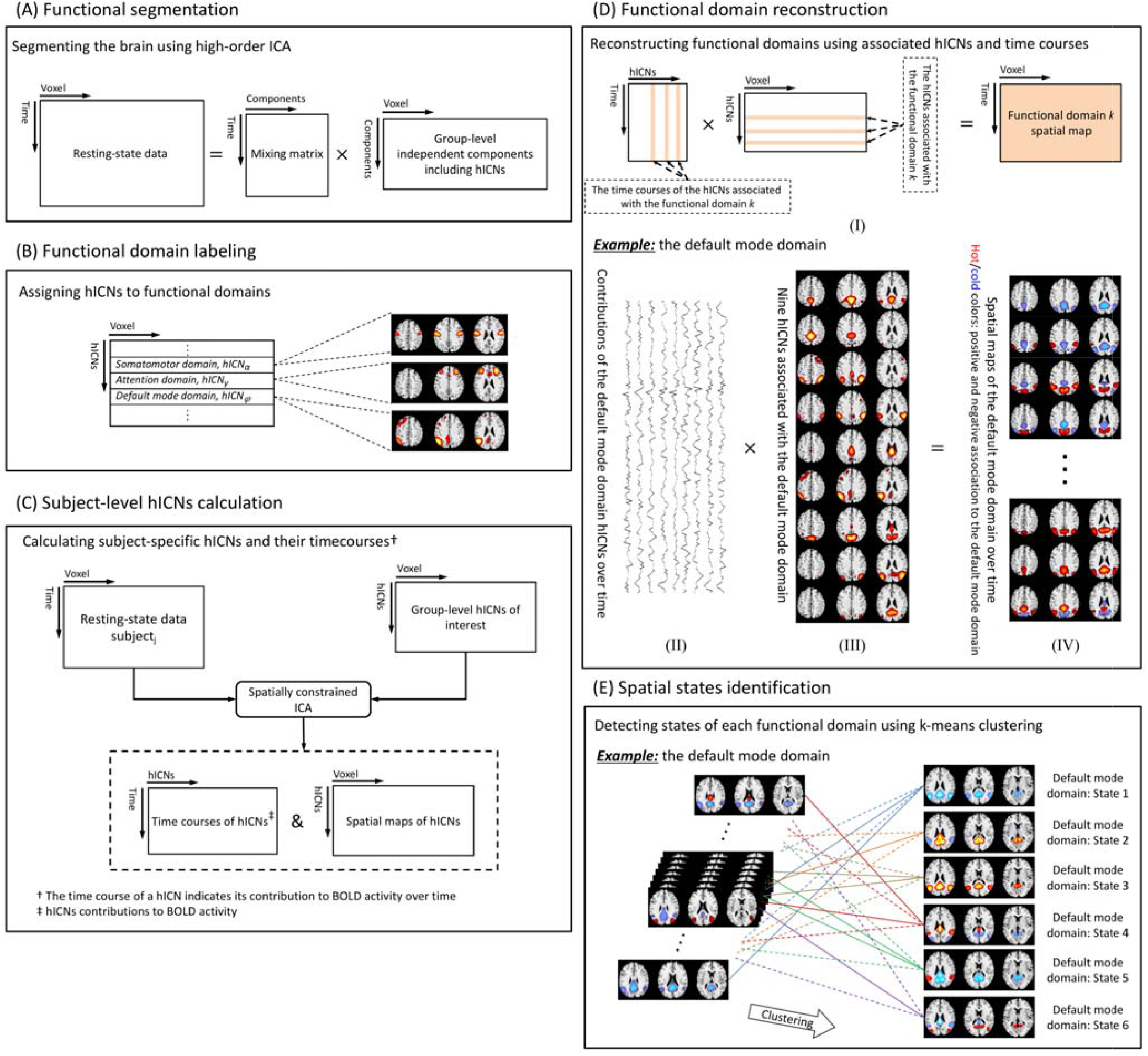
Schematic of the analysis pipeline. (A) High-order group-level spatial ICA (# Components = 200) was applied on the proce resting-state fMRI data from 309 individuals. (B) Sixty-five components were identified as the high-order intrinsic connectivity netw (hICNs) of interests and assigned to one of nine cortical and subcortical functional domains (FDs) including attention, auditory, de mode, frontal default mode, frontoparietal, language, somatomotor, subcortical, and visual domains (Figure 3). (C) Spatially constra ICA (Lin et al., 2010) was used to estimate the time courses of hICNs and their spatial maps for each individual. The time course of a indicates the contribution of the hICN at different time points. (D) FDs were reconstructed using the linear combination of the assoc hICNs and their contributions at any given time. (E) Spatial domain states associated with each FD were estimated using k-m clustering on the spatial maps of the FD.

### 2.4. Functional Domain (FD) Construction

At any given time point, FDs were constructed from the associated hICNs and their contributions as follows. First, subject-specific hICNs and their time courses were calculated via a spatially constrained ICA approach using the group-level hICNs as references (Figure 2.C) (Lin et al., 2010). The time course of each hICN describes its temporal evolution and represents its contributions to the BOLD signal over time. Next, to reduce noise, a post-hoc cleaning procedure was also performed. For this purpose, various cleaning procedures were compared. The evaluated post-hoc cleaning procedures include (C1) orthogonalizing with respect to estimated subject motion parameters, linear detrending, despiking, and band-pass filtering using a fifth-order Butterworth (0.001-0.15 Hz), (C2) replacing the Butterworth filter with a Gaussian moving average filter with different window sizes (10×TR to 90×TR) and keeping the rest of cleaning steps the same as the first procedure, (C3) using only a Gaussian moving average (with different window size from 10×TR to 90×TR), and (C4) no post-hoc cleaning procedure. The cleaning procedures were also evaluated for two scenarios: 1) cleaning procedures were applied on the time courses of hICNs, and 2) cleaning procedures were applied on voxel-level after reconstructing the spatial maps of FDs. The various approaches resulted in almost identical spatial domain states (the definition of spatial domain states in *Section 2.5*) suggesting that post-hoc cleaning procedures do not substantially alter the dynamic properties of FDs. Given the similarity of the two scenarios, we suggest utilizing the first and applying cleaning procedures on the time courses of hICNs as the computational load is much lower. C1 procedure was selected a post-hoc cleaning procedure, as it is commonly used as a cleaning procedure and has previously demonstrated its effectiveness at noise reduction (Damaraju et al., 2014). Finally, after the post-hoc cleaning step, each FD was reconstructed using the linear combination of the associated hICNs and their contributions at any given time point resulting in 49749 (309 subjects × 161 time points) spatial maps for each FD (Figure 2.D and Equation.1).

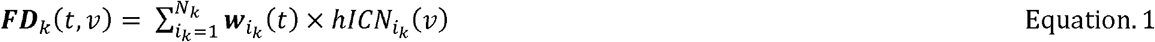

Where ***FD*** *_k_*(*t,v*), is the FD *k* at the time point *t,v* is voxel index, N*_k_* is the number of hICNs belongs to the FD *k*, 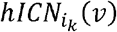 is the hICN 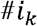 of the FD*k*, and 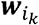(*t*)is the contribution of 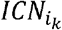 at the time point t.

### 2.5. Spatial Domain States Identification and Verification

The spatial dynamics of FDs were captured via spatial domain states. The spatial domain states for a given FD are a set of distinct spatial patterns and can be obtained using a clustering approach (Figure 2.E). Here, we used k-means clustering, and the correlation distance metric was used as the distance function because it detects spatial patterns irrespective of voxels’ intensities. The number of states (clusters) for each FD was determined using the elbow criterion by searching for the number of clusters from 3 to 15 (Damaraju et al., 2014; Yaesoubi et al., 2017). Similar to what we have done previously (Allen et al., 2014), initial clustering was performed on a subset of the data exhibiting maximal deviation from the mean (called exemplars) and was repeated 100 times with different initializations using k-means++ (Arthur and Vassilvitskii, 2007). Exemplars are the data points in which the amount of variance explained by either of hICNs is significantly (p < 0.001) higher than the average amount of variance explained by hICNs across the whole dataset (49749 fMRI volumes). The estimated centroids from initial clustering using exemplar were then used as cluster center initializations to cluster the whole dataset.

We further verified the spatial domain states by evaluating the average BOLD signal of associated regions across states. In other words, we examine how well variations in the FDs reflect the underlying BOLD signal. Let us assume region *j* is only associated with FD *k*. Then, if the association of region *j* to FD *k* is positive/negative at state *i*, the neural activity of region *j* measured by the BOLD signal at the state *i* of FD *k* should be above/below its average (i.e. the average BOLD signal of region *j*). We expect to observe a very similar pattern of agreement between the regions’ associations to FDs and their amplitude of BOLD signals even if regions are simultaneously involved in different FDs at a given spatial domain state.

### 2.6. Spatial Dynamic Evaluation

To study the spatial dynamics of FDs, we first evaluated variations in regions’ associations with each FD across different spatial domain states. A voxel-wise, one sample t-test was applied to the data of each state (i.e. the spatial maps of a given FD at time points belongs to the spatial domain state), and the average *t*-value was calculated for 246 regions of the Brainnetome atlas (Fan et al., 2016). A region was assigned to a spatial domain state if its average *t*-value falls outside the Tukey inner Fences (below lower inner fence or above upper fence). A Tukey inner fence is defined as [Q_1_-1.5× (Q_3_-Q_1_) Q_3_+1.5× (Q_3_-Q_1_)], where Q_1_ and Q_3_ are the first and third quartiles (Hoaglin et al., 1986). Next, we investigated the overall spatiotemporal variations within each FD. Previously, Cole et al. developed an index called the “global variability coefficient (GVC)” to evaluate variations in the connectivity of the brain networks across different tasks using multi-task fMRI data (Cole et al., 2013). Here, we introduced a related measure called the variability index (VI) to evaluate the level of variability for each FD. Similar to GVC, VI is defined as the standard deviation of a region’s association to an FD which can be estimated using the standard deviation equation of binomial distribution. For example, if FD *i* has 5 states, and region *j* is involved in only one state, the standard deviation of the region *j* being associated with the FD *i* is (5 × 0.2 × 0.8)^0.5^. The average of VI values within each FD characterizes the overall spatiotemporal variability of the FD.

### 2.7. Functional State Connectivity and Functional Modules (FMs)

Like other structures of this hierarchical functional architecture, FDs interact with each other. To evaluate these interactions, we need to calculate functional connectivity between FDs. Functional connectivity is defined as the temporal dependency of neural activity (Friston et al., 1993). In fMRI, functional connectivity is typically measured by calculating the temporal coherence between BOLD time series or time series associated with brain networks. Using the same strategy, we can estimate functional connectivity between FDs by calculating the temporal coherence (coupling) between states of FDs. For this purpose, we calculate the level of concurrency between pairs of states using a coincidence index known as the Dice similarity coefficient (DSC) (Dice, 1945). The functional inter-domain state connectivity (called functional state connectivity for clarity) is defined as the ratio between the number of time points in which two states from different FDs occur simultaneously and the average occurrence of both states. In other words, functional state connectivity between state *i* of FD *m* (*FD* _*m,i*_) and state *j* of FD *n* (*FD* _*n,j*_) was calculated as the ratio of the number of the time points that state *i* of FD *m* and state *j* of FD *n* occurred simultaneously to the average occurrence of state *i* of FD *m* and state *j* of FD *n* (Equation. 2).

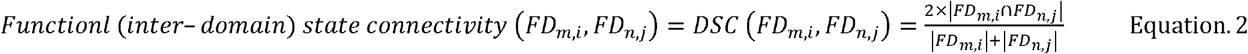

We further used functional state connectivity values to identify FMs. A FM is defined as a set of spatial domain states of FDs that reoccur together frequently in a distinguishable manner. In other words, a set of spatial domain states with higher connectivity with each other than with other states. FMs can be extracted using graph-based community detection approaches like the Newman modularity detection approach (Newman, 2006).

### 2.8. Group Comparison Analysis

The clinical utility of our approach was evaluated by comparing spatial domain states of FDs between patients with SZ and healthy controls. For each region associated with a given spatial domain state, the average value of the FD was compared between patients with SZ and healthy controls using a general linear model (GLM) with age, gender, data acquisition site, and mean framewise displacement (Power et al., 2012) as covariates. Framewise displacement measures instantaneous head motion as a single scalar value by calculating changes in the six rigid body transform parameters (framewise displacement(t) = |Δdx(t)| + |Δdy(t)| + |Δdz(t)| + |Δα(t)| + |Δβ(t)| + |Δγ(t)|) and was included as a covariate to mitigate effects of head motion (Power et al., 2012). Statistical comparison was further performed on FMs by comparing each pair of functional state connectivity between patients with SZ and healthy controls using the same procedure explained in section 2.8. For all analysis, statistical results were corrected for multiple comparisons using a 5% false discovery rate (FDR) (Benjamini and Hochberg, 1995).

## 3. Results

### 3.1. High-order Intrinsic Connectivity Network (hICN) Extraction

Figure 3 displays the composite view of the hICNs selected from the group-level spatial ICA results. Among 200 independent components, sixty-five were selected as cortical and subcortical hICNs and categorized into nine FDs. The nine FDs were defined based on the prior knowledge from previous studies (Allen et al., 2014; Allen et al., 2011; Damaraju et al., 2014; Iraji et al., 2016) and large-scale brain networks obtained from low-order ICA. The nine FDs are attention (Allen et al., 2011; Damoiseaux et al., 2006; Lee et al., 2013), auditory (Allen et al., 2014; Allen et al., 2011; Damoiseaux et al., 2006), default mode (Allen et al., 2011; Iraji et al., 2016; Zuo et al., 2010), frontal default mode (Iraji et al., 2016; Zuo et al., 2010), frontoparietal (Allen et al., 2011; Iraji et al., 2016; Lee et al., 2013; Zuo et al., 2010), language (Lee et al., 2013; Tie et al., 2014), somatomotor (Allen et al., 2011; Damoiseaux et al., 2006; Iraji et al., 2016), subcortical (Allen et al., 2014; Allen et al., 2011), and visual (Allen et al., 2011; Damoiseaux et al., 2006; Iraji et al., 2016; Zuo et al., 2010). hICN selection and FD labeling were performed using the anatomical and presumed functional properties of hICNs, and their relationships with large-scale brain networks obtained from low-order ICA. Detailed information of the hICNs including spatial maps, coordinates of peak activations, and temporal information can be found in Supplementary 1. The selected hICNs are primarily in cortical and subcortical gray matter and show high spatial similarity with hICNs identified in pervious works (Allen et al., 2011; Damaraju et al., 2014). After selecting hICNs of interest, spatially constrained ICA was utilized to calculate the subject-specific hICNs and their time courses (Figure 2.C) which were further used to reconstruct the FDs for each individual and time point.

**Figure 3.**
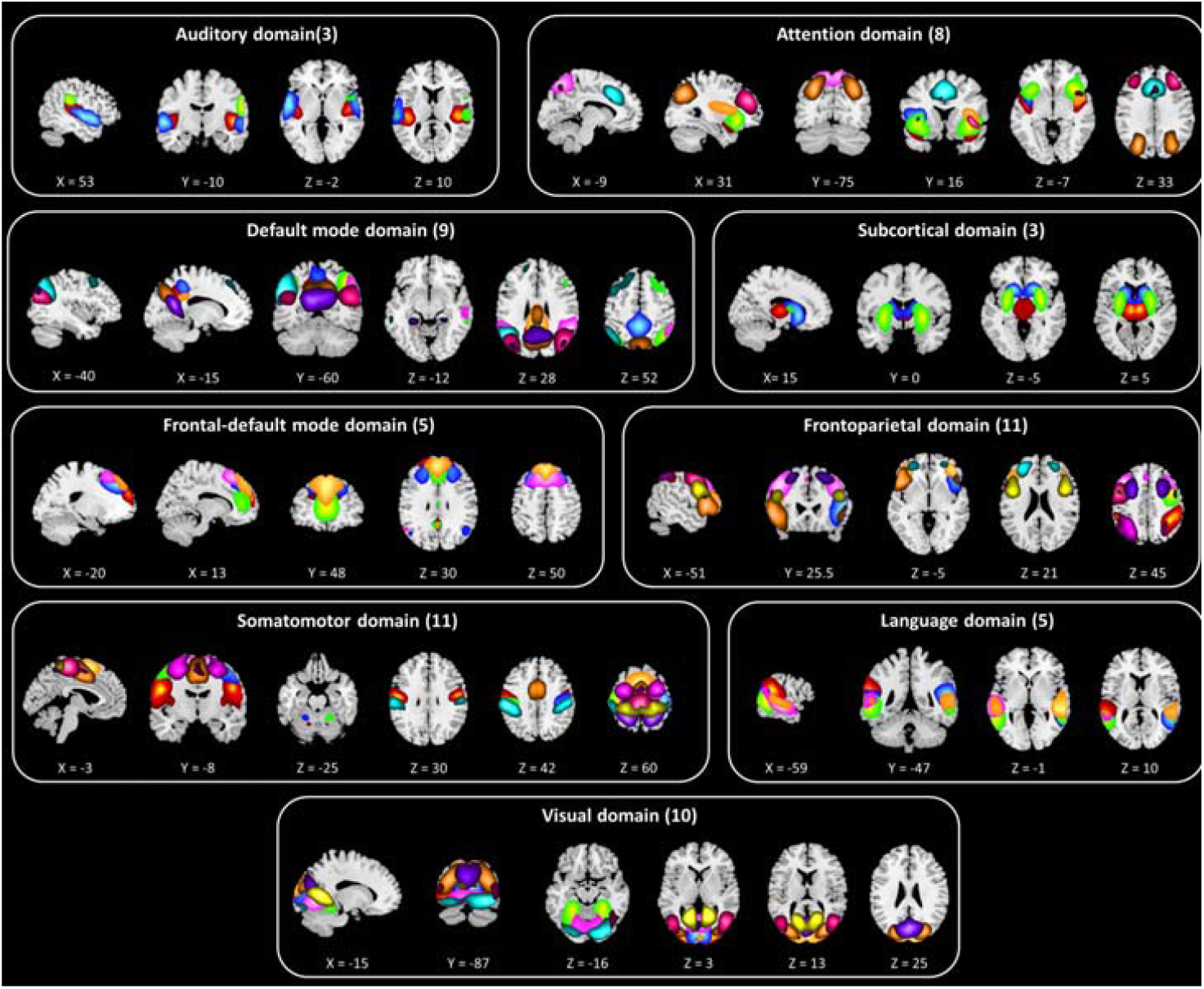
Composite maps of nine functional domains (FDs) generated from the sixty-five high-order intrinsic connectivity networks (hICNs). Each color in a composite map corresponds to one of hICNs associated with the given FD. The detailed information of hICNs can be found at Supplementary 1.

### 3.2. Functional Domain Construction

At each time point, FDs were reconstructed using the associated hICNs and their contributions at that time point, resulting in 49749 (309 subjects × 161 time points) spatial maps for each FD. Figure 2.D illustrates an example of FD reconstruction for the default mode domain which contains nine hICNs shown in Figure 2.D(III). At each time point, the default mode domain (Figure 2.D(IV)) was calculated using the linear combination of the nine hICNs, and their contributions (Figure 2.D(I)). In Figure 2.D(IV), **hot** and **cold** colors represent positive and negative associations to the default mode domain. The spatial maps of FDs over time for randomly selected individuals are provided as Supplementary movies 1–9 and also at the following link https://www.youtube.com/playlist?list=PLZZPPK0O_qFuil41n4U_HSZ668cFDuG7l. FDs display highly dynamic behaviorss, and the contributions of brain regions to FDs vary significantly over time. Brain regions show both strong positive and strong negative associations to FDs over time. Moreover, as will be demonstrated later in *Section 3.4*., variations in regional association to FDs go beyond amplitude modulation. For example, some brain regions which are strongly involved in FDs at a given time point become dissociated at other time points.

### 3.3. Spatial Domain State Identification and Verification

K-means clustering was applied to the spatial maps of each FD and summarized into a set of reoccurring spatial patterns called spatial domain states. Figure 4 and Supplementary 2 show the spatial maps of cluster centroids as representations of spatial domain states. The number of clusters (*k*) for each FD was determined using the elbow criterion. Additionally, exploratory analyses over a large range of *k* demonstrates that these clusters are fully reproducible and spatial domain states are very similar (see Supplementary 3). In general, we can categorize the states into voxel-wise coherent and incoherent states. In voxel-wise coherent states, the regions associated with FDs show a similar pattern of association, either positive or negative, while voxel-wise incoherent states contain regions with both positive and negative associations to FDs. In Figure 4 and Supplementary 2, the total number and percentage of states occurrences are listed above each centroid. Occurrence rates range from 10% to 25%. For all FDs, the top two dominate states are voxel-wise coherent states with occurrence rates above 20%.

**Figure 4.**
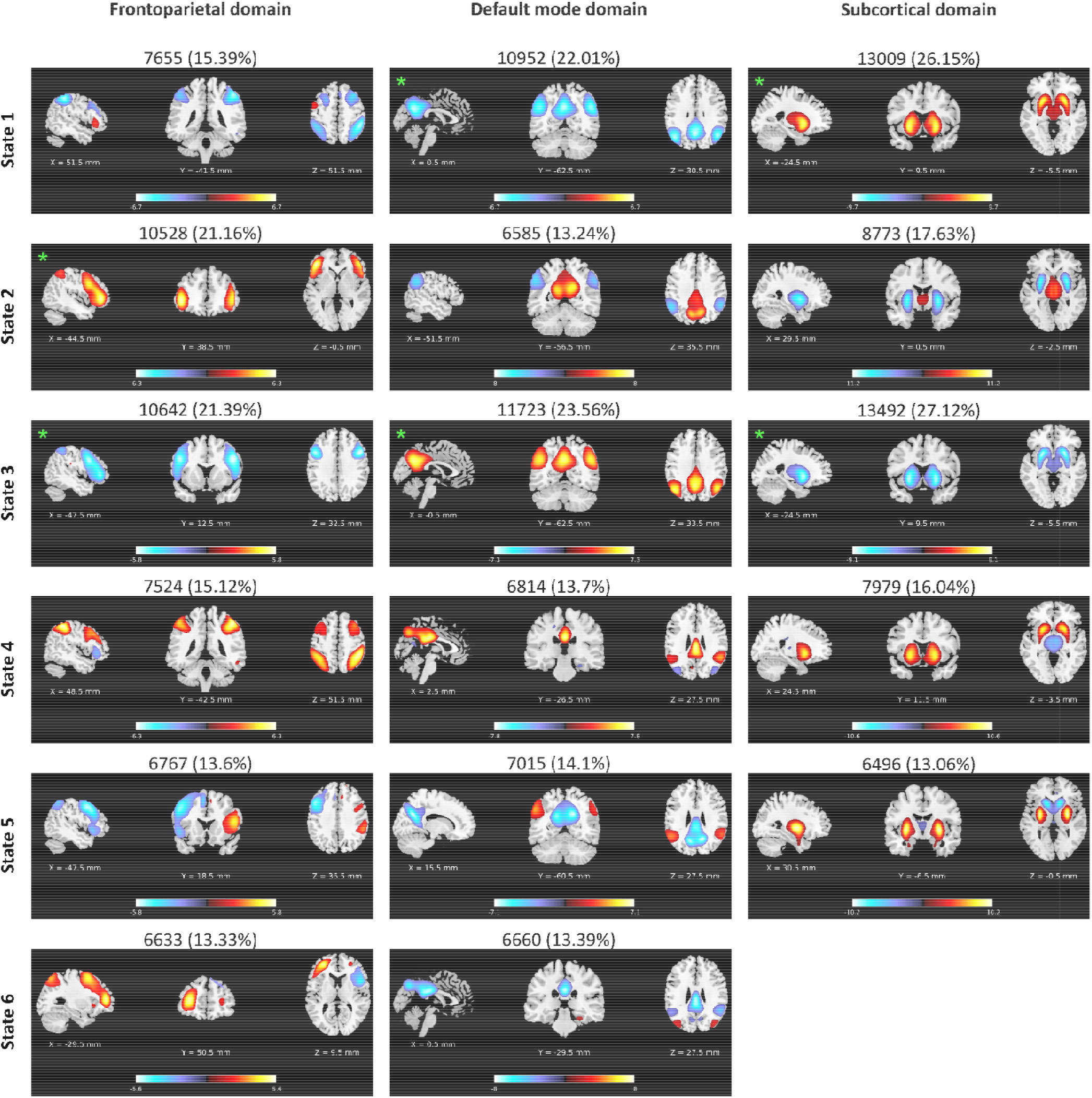
Examples of spatial domain states associated with three functional domains (FDs) including frontoparietal, default mode subcortical domains. Orthogonal views of spatial domain states, thresholded at |Z| > 1.96 (*p* = 0.05). Each spatial map represent centroid of a cluster; and sagittal, coronal, and axial slices are shown at the peak activation of the centroids. Hot and cold colors repr positive and negative association of voxels to the FDs. The total number and percentage of occurrences are listed above each cent Voxel-wise coherent states were marked using green asterisks. In voxel-wise coherent states, the associated regions show a similar pa of association (either positive or negative), and voxel-wise incoherent states contain both regions with positive and negative associatio FDs. The spatial domain states for all nine FDs can be found at Supplementary 2. For further details regarding variations in reg associations to FDs please see Figure 6.

Investigating the relationships between BOLD signal of regions and their contributions to FDs exhibit overall the same pattern that regions have higher/lower activity when they have positive/negative association to their corresponding FDs. We observe this agreement between BOLD signals and regions associations to FDs for 96.02% of the cases. An example of the relationship between BOLD signal and regional association is presen in Figure 5 and Supplementary 4. Further investigation determined that different directionalities between regi associations to FDs and the amplitude of their BOLD signals only occurs in regions with weak contributio FDs and/or small BOLD signal difference from the baseline (see Supplementary 5).

**Figure 5.**
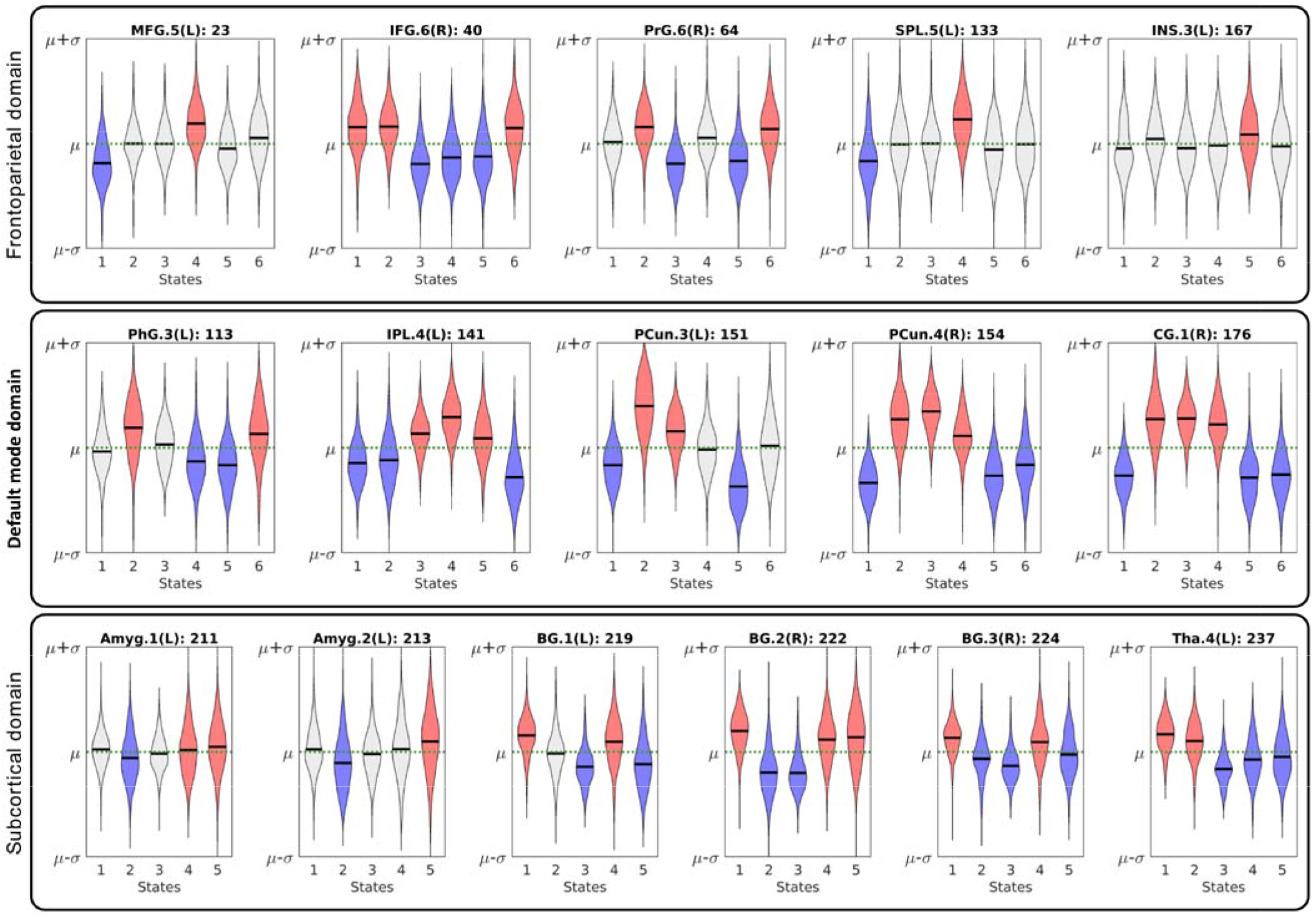
The relationship between the amplitude of the BOLD signal and regions associations with functional domains (FDs). Exampl average BOLD signal of regions across spatial domain states for the same three FDs as Figure 4. Example for all nine FDs can be fou Supplementary 4. Each violin plot represents the average BOLD signal of a region for spatial domain states of a given FD. The μ (g dashed line) and σ are the average and standard deviation of BOLD signal for each region across all time points, and the black represents the average of the BOLD signal of a region across all subjects in the corresponding state. Red and blue colors represent the states with positive and negative associations of regions to FDs, the light gray color indicates no association between corresponding FD region. The results suggest regions associations to FDs are related to their neural activities measured by the amplitude of BOLD signal. When regions are positively/negatively associated with FDs, their average BOLD signal are above/below their own average across all time points. This suggests the variations in a region’s association to an FD is related to its neural activity, as observed by the amplitude of BOLD signal. The abbreviation and regions labels listed above each violin plot are based on the Brainnetome atlas (Fan et al., 2016). Amyg (Amygdala), BG (Basal Ganglia), CG (Cingulate Gyrus), IFG (Inferior Frontal Gyrus), INS (Insular Gyrus), IPL (Inferior Parietal Lobule), MFG (Middle Frontal Gyrus), PCun (Precuneus), PhG (Parahippocampal Gyrus), PrG (Precentral Gyrus), SPL (Superior Parietal Lobule), and Tha (Thalamus).

### 3.4. Spatial Dynamic Evaluation

Our analysis reveals that FDs are spatially fluid, and brain regions are transiently associated with FDs. FDs display distinct spatial patterns across their spatial states which include changes in the regions associated with them. Figure 6 and Supplementary 6 summarize variations in regions associations to FDs in which *t*-value indicates the strength of each association. The results highlight changes in regions’ memberships and the strengths of their associations to FDs over time. As an example, CG4 (cingulate gyrus subregion 4) is positively associated with the default mode in State 2 and 3 (presented in **hot** color), negatively associated with the default mode at States 1 and 5 (presented in **cold**), and becomes dissociated at States 4 and 6 (presented in gray). The list of associated regions, their coordinates, and the strengths of their associations to FDs can be found at Supplementary 7. The variations in regions membership to FDs over time can potentially explain inconsistencies in findings of previous static analyses regarding regions memberships to brain networks. Furthermore, examining the overall spatiotemporal variations of FDs using a variability index (VI) reveals that the FDs such as the frontoparietal and attention which are engaged in a wide variety of cognitive functions have higher variations than other FDs. The mean and standard error of VI values are listed above the chart of each functional domain in Figure 6. Interestingly, in a previous multi-task fMRI study, the same patterns variations was observed across a variety of tasks in which frontoparietal, attention and auditory, in order, show the highest variation, and the subcortical has the lowest changes in their connectivity patterns (Cole et al., 2013).

**Figure 6.**
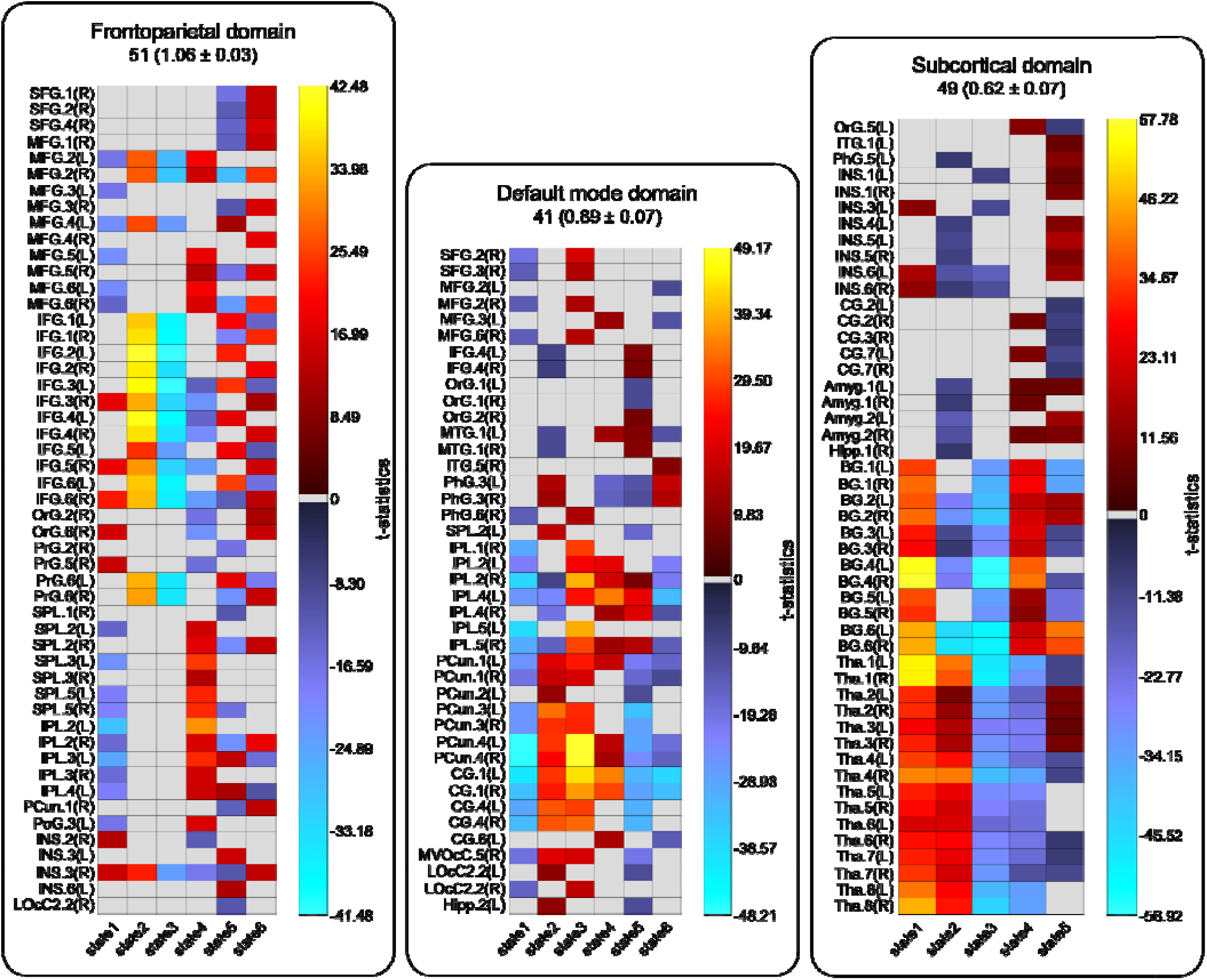
Spatiotemporal variations of functional domains (FDs). The chart represents the regions associated with the frontoparietal, default mode, and subcortical domains. Spatiotemporal variations of all nine FDs can be found at Supplementary 6. The brain anatomical parcellation is based on the Brainnetome atlas. The total number associated regions and the mean and standard error of variability index(VI) are listed above each chart. The results show different regions are associated to FDs at different states. **Hot** and **cold** colors represent positive and negative associations and **gray** represents dissociation of the regions at the states. *t*-value indicates the strength of the regions association to FDs. VIs represent the overall variability of each FD where the frontoparietal and subcortical domains shows maximum and minimum variations. The abbreviation and regions labels are the same as defined in the Brainnetome atlas. SFG (Superior Frontal Gyrus. MFG (Middle Frontal Gyrus), IFG (Inferior Frontal Gyrus), OrG (Orbital Gyrus), PrG (Precentral Gyrus), MTG (Middle Temporal Gyrus. ITG (Inferior Temporal Gyrus), PhG (Parahippocampal Gyrus), SPL (Superior Parietal Lobule), IPL (Inferior Parietal Lobule), PCum (Precuneus), PoG (Postcentral Gyrus), INS (Insular Gyrus), CG (Cingulate Gyrus), MVOcC (MedioVentral Occipital Cortex), LOcC (lateral Occipital Cortex), Amyg (Amygdala), Hipp (Hippocampus), BG (Basal Ganglia), and Tha (Thalamus).

### 3.5 Group Differences in Spatial Domain States

The spatial domain states of FDs were compared between patients with SZ and healthy controls using a regression model including age, site, gender, and mean framewise displacement as covariates. Several FDs reveal significantly weaker activity across different states in patients with SZ compared to healthy controls. In general, patients with SZ showed reductions in the regions’ dynamic associations with FDs, except the dorsolateral region of Brodmann area (BA) 37 in State 5 of the language domain. Among all FDs, the visual, subcortical and attention domains are most affected. For the visual domains, all states except State 5 show significant differences between two groups. The most affected regions in the visual domain include the ventromedial occipital cortex, the lateral occipital cortex, and the ventromedial fusiform gyrus. In the subcortical domain, the thalamus showed significant differences in States 1, 2, and 3. For the attention domain, the left insula and the opercular area of left BA 44 in States 1 and 2, and the lateral area of BA 38 in State 4 show the highest differences. The other regions with significant differences in the language domain including right BAs 41/42 and the rostral area of left BA 22 demonstrated decreased associations in patients with SZ similar to the general pattern. Exploratory analyses on a subsample of the data with little head motion and no significant difference in mean framewise displacement between two groups (*p* = 0.5) displayed a similar pattern in group differences as using the full dataset. The full details of spatial comparison and regions with statistical differences can be found in Supplementary 8. Finally, in an exploratory analysis, we calculated the average spatial maps of FDs over time and compared them between two groups, healthy controls and patients with SZ. No significant difference was observed between the two groups. Furthermore, we have also compared the static spatial independent components obtained from low-order ICA and only observed differences between independent components associated with the subcortical, language, and attention domains, thus missing most of the changes identified in this work due to the assumption of spatially static networks. This highlights the importance of using spatial dynamics to examine the spatial patterns of FDs.

### 3.6 Functional State Connectivity and Functional Modules

The functional state connectivity matrix was estimated by calculating the temporal coupling between the states of FDs using DSC index (Figure 7). Using the Newman modularity detection approach, seven FMs were detected (Figure 8). Investigating group differences reveals an overall decrease in functional state connectivity within FMs in patients with SZ (the **green** lines in Figure 8). Differences between patients with SZ and healthy controls are the most pronounced in FM 1 which mainly includes hypoconnectivity between the subcortical and others domains (Figure 8). Hypoconnectivity of the subcortical domain, which is also observed in other FMs, is the largest patient/control differences between groups. Note that the subcortical domain demonstrates alterations in both its activity patterns and its connectivity with other FDs in States 1, 2, and 3, which suggests it was the major source of the observed hypofunction in schizophrenia. Alteration in functional state connectivity of the default mode within FMs is another interesting finding. Although the comparisons of the spatial domain states of the default mode between two groups did not reveal any significant difference (3.5. Group Differences in Spatial Domain States), hypoconnectivity between the default mode and several FDs was observed within FMs. Interestingly, we observed hypoconnectivity between State 2 of the default mode domain and State 5 of the auditory domain, even though neither show significant change in their activity patterns in patients with SZ. This suggests that alterations in functional connectivity can occur in the absence of change in functional activity and vice versa.

**Figure 7.**
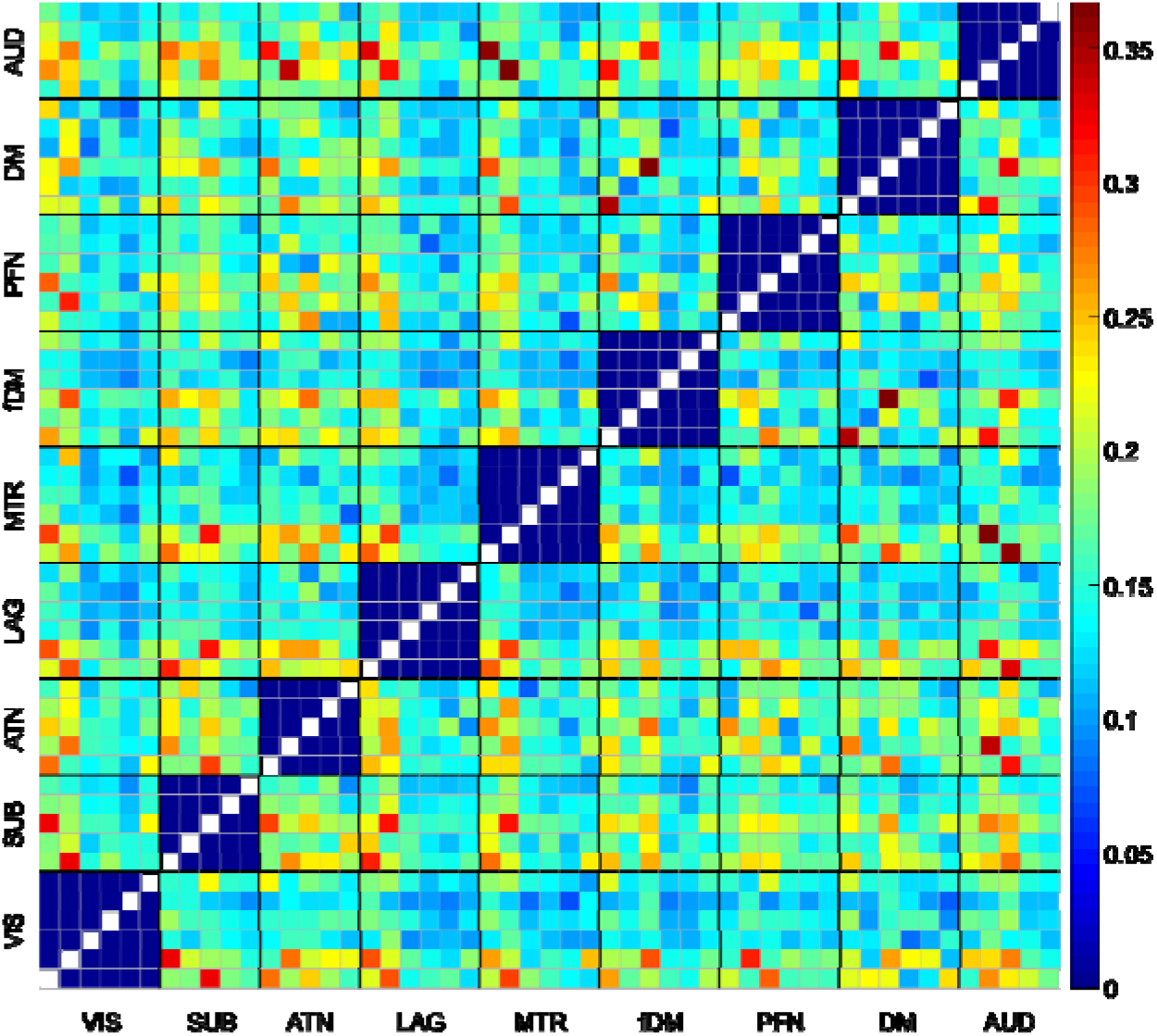
Functional state connectivity was estimated by calculating the level of concurrency between spatial domain states using DSC index. VIS (visual domain), SUB (subcortical domain), ATN (attention domain), LAG (language domain), MTR (somatomotor domain), fMD (frontal default mode domain), PFN (frontoparietal domain), DM (default mode domain), and AUD (auditory domain).

**Figure 8.**
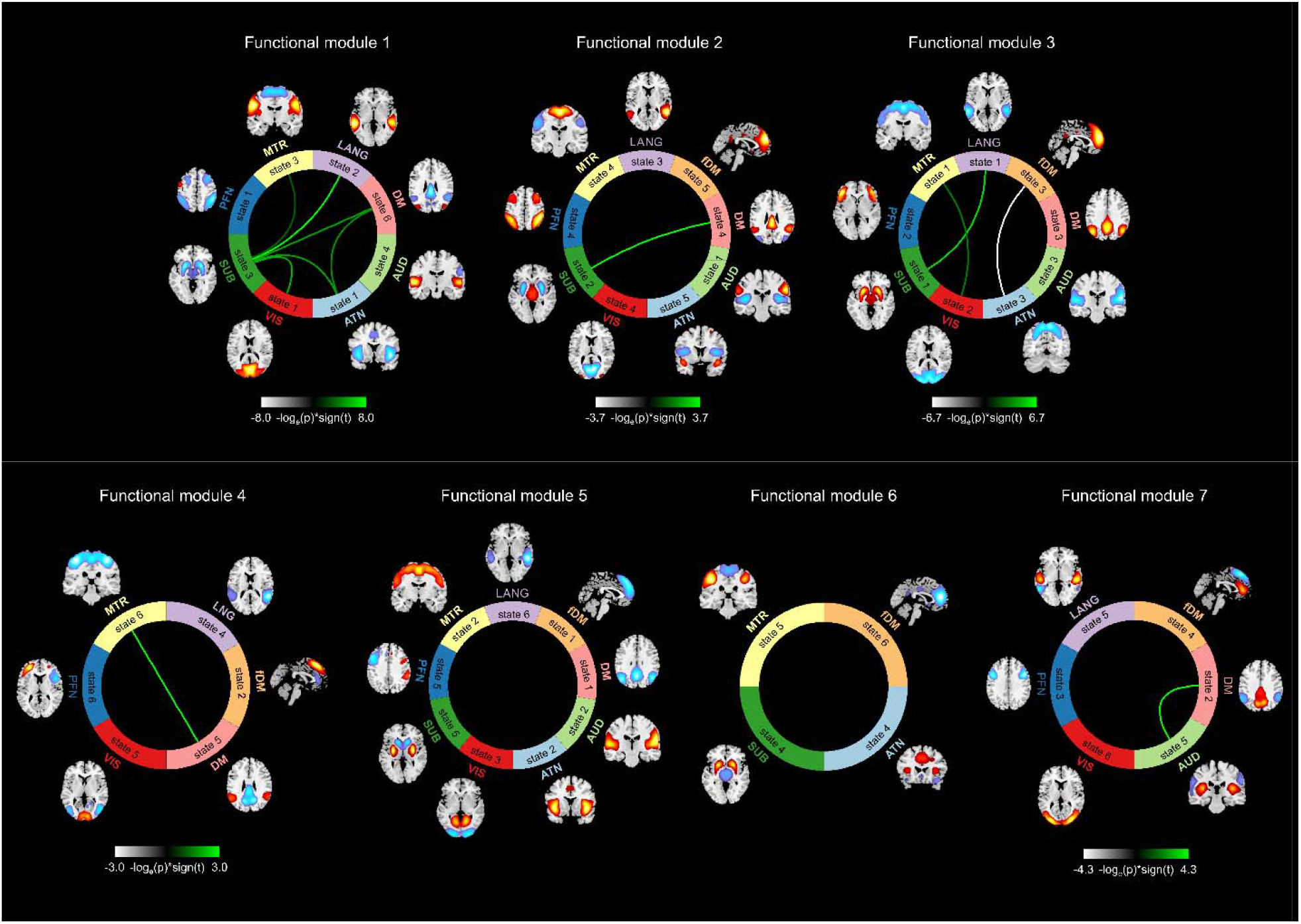
Functional modules (FMs) and functional state connectivity comparisons between healthy controls and patients with schizophrenia (SZ). Seven FMs were detected using Newman modularity detection approach. The **green** lines represent higher functional state connectivity in healthy controls than patients with SZ, and the **silver** one shows a higher functional state connectivity in patient with SZ.

## Discussion

The brain reorganizes itself on different temporal and spatial scales which manifest at the macroscaleas variations in both temporal and spatial couplings of brain functional organizations. Recent findings display the abiliy of fMRI to capture time-varying information of the brain (Calhoun et al., 2014; Hutchison et al., 2013; Preti et al.,2017). However, the majority of these studies have overlooked the spatial variations of brain functional organizations. In the present study, we propose an approach that captures the spatiotemporal variations of FDs, i.e., spatial dynamics, using the brain functional hierarchy model at macroscale. In agreement with our hypothesis, we observed that FDs are evolving spatially over time. Evaluating the spatial dynamics of individual FDs revealed a set of distinct, reoccurring spatial patterns (spatial domain states) within each FD. Variations in the spatial patterns of FDs over time were further accentuated by changes in regions’ memberships to FDs. For example, brain regions join FDs and dissociate from them over time. In early work, Cole et al. used multi-task fMRI data and predefined anatomical regions and showed changes in the spatial patterns of brain networks across various task scenarios (Cole et al., 2013). Here, we demonstrate that spatial variations exist even in a resting state of the brain due to the dynamic nature of the brain. Interestingly, we observe similar results to task data in which the frontoparietal, attention and auditory domain, in order, show the highest variation, and the subcortical has the lowest changes in its connectivity pattern. Spatial dynamics may also explain the inconsistencies observed in previous static analyses regarding regions’ memberships to FDs (also known as large-scale brain networks). For example, previous static analysis reported different sets of regions for each brain network, including different sets of regions for the default mode (Andrews-Hanna et al., 2010; Buckner et al., 2008; Damoiseaux et al., 2006; Fox et al., 2005; Garrity et al., 2007; Greicius et al., 2003; Lee and Xue, 2018; Shirer et al., 2012; Wang et al., 2014; Zuo et al., 2010). We suggest that different sets of regions are associated with a given FD at different time points, and only an overall dominant pattern is identified in the static analysis. In our opinion, alternative interpretations for the observed inconsistency in the spatial patterns of FDs can be the interdigitated parallel networks observed in a previous single-subject study (Braga and Buckner, 2017). Their findings suggest that each brain network may consist of parallel networks that work simultaneously, and only one or combination of them is captured in group-level analysis. The concept of the spatial dynamic is not against the existence of interdigitated networks within each FD, as they can be extracted using the time points during which they contribute the most in the spatial patterns of the given FD. For instance, Braga and Buckner, 2017, observed two district patterns for the default mode. One includes the parahippocampal cortex and posterior inferior parietal lobule, while the other includes anterior inferior parietal lobule regions. We found similar variations in the parahippocampal gyrus and inferior parietal lobule, along with many other regions across default mode states.

The concept of spatial dynamics and variations in regions’ associations over time was also confirmed via the amplitude of BOLD signal. We observed a direct relationship between the activities of regions measured by the BOLD signal and their contributions to FDs which supports our proposition that regions have higher/lower activity than their baseline when they have positive/negative association to their corresponding FDs. It should be noted that despite the high agreement between a region’s association to FDs and its BOLD signal, the spatial dynamics of FDs cannot be captured by directly applying a clustering approach to the BOLD signal. (High-order) ICA enables us to parcellate the brain to the functional units from the data itself to assure functional homogeneity, something which is not provided when using predefined regions from atlases (Yu et al., 2017). Moreover, using predefined spatial nodes instead of hICNs ignores simultaneous roles for brain regions. Using predefined atlases instead of hICNs also limits our ability to detect the spatial variations of FDs over time as the variations would become limited to sets of predefined regions. Most importantly, the goal of this study is to capture the spatial dynamics of FDs, which cannot be achieved by directly using the BOLD signal of a set of predefined regions as the unprocessed BOLD signal do not convey information regarding their contributions to a given FD. Therefore, using the BOLD signal directly only measures variations in activity patterns of regions over time rather than spatiotemporal variations of FDs.

### Hierarchical Approach: Strengths, Limitations, and Future Directions

Our proposed approach to capture the spatiotemporal variations of FDs is based on the well-accepted assumption that the brain can be modeled as a hierarchical functional architecture with different levels of granularity. Each level of this architecture includes several elements, with each element involved in specific functions, and the elements of higher levels have less functional homogeneity and increased dynamic behavior. Constructing functional hierarchy requires identifying functional units, which is quantified as a pattern of regions with the same functional activity over time (high functional homogeneity) and can be extracted from the data. We suggest hICNs are good approximations for functional units. Estimation of functional units and spatially fluid properties are limited by data quality, such as spatial and temporal resolutions, and the properties of the imaging modalities. Although one advantage of the proposed approach is capturing the spatiotemporal variations of FDs up to the maximum temporal and spatial resolutions exist in the data, we can improve the functional granularity and reconstruct the hierarchy from lower levels, such as cortical columns, by adjusting the data acquisition and analytical approaches. We can also use other imaging modalities, such as calcium imaging (Matsui et al., 2018) or photoacoustic tomography (Nasiriavanaki et al., 2014), to estimate functional units and construct functional hierarchy, which can help to improve our understanding of the spatially fluid properties and information processing. Another advantage of the approach is employing both spatial and temporal information of ICNs rather than only using either ICNs’ spatial patterns or their time courses as it is the common strategy in previous work.

Moreover, our approach provides information about each FD at every time point. Because at any given time point, FDs are constructed from hICNs’ contributions of the same time point in contrast to being influenced by other time points, the approach can capture the spatial dynamic regardless of temporal resolution. In fMRI, this means the approach has the ability to detect dynamic patterns independent of TR. Of course, having data with high temporal resolutions and/or from a longer periods of time can potentially provide us more information regarding dynamic behavior of FDs including the detecting new dynamic patterns. In addition to spatial dynamics of FDs, we also demonstrate the ability of the approach in capturing time varying properties at a global scale. For this purpose, we developed an index called functional (inter-domain) state connectivity to compute temporal coupling between FDs and calculate FMs. Future studies are required to capture spatial and temporal dynamic properties within and between different levels of hierarchy. As the first step, a future study should investigate the temporal properties of spatial domains states and associated dynamic indices such as dwell time, leave time, fraction rate, and transition matrix. It is apropos to mention that, in this study, k-means clustering was used as a tool to examine variations existing in brain functional domains, but other methods are equally applicable. While k-means clustering yields valuable results, neither the states identified using k-means clustering are likely to be the true origins of time-varying behavior of FDs, nor the assumption that there is only activity state per time point, as activity patterns of a given FD occur simultaneously. Further studies are therefore needed to find improved representations of the time-varying behavior of FDs.

The most challenging step of the approach requiring future investigation and further improvement is the estimation of functional units and their FD labeling. Although our approach allows the detection of spatial variations over time regardless of a selected partitioning procedure (i.e. selecting functional units and associated FDs), good partitioning is essential to fully capture the dynamic characteristics of the data. High-order ICA has several advantages over predefined atlases including: 1) high-order ICA allows the segregation of functional roles of individual regions; 2) each hICN is a pattern of functionally homogeneous regions extracted from data itself, which is closer with the definition of functional units than predefined anatomical regions; and 3) the spatial variations of high-order ICA are not limited to fixed regions as with predefined atlases and allow for individual variability in the spatial maps (Allen et al., 2012; Calhoun and Adali, 2012). However, despite the advantages of hICNs, the level of parcellation (i.e. a number of components) requires further investigation. In addition to the number of hICNs, hICN grouping (i.e. FD labeling) is another key piece of the hierarchical approach. In our current work, hICNs were grouped into nine FDs using a semi-automatic process approach including 1) our prior knowledge from anatomical and functional properties; 2) the relationship of hICNs with results of low-order ICA; and 3) similarity between their timecourses of hICNs. While different grouping approaches are acceptable, a robust, data-driven approach is recommended for FD identification and hICN assignment. Furthermore, changes in hICNs’ memberships to FDs over time is another crucial factor that needs to be considered. The present study limits assigning each hICN to one FD; however, hICNs can also change their memberships to different FDs over time. Ongoing work is assigning hICNs to FDs at any given time point using the information of the data in that time point.

### Schizophrenia

Schizophrenia is a heterogeneous disorder characterized by symptoms of impaired reality testing such as hallucinations, delusions, and frequently disorganized speech and behavior, as well as impairments in cognition across a range of domains (Association, 2013). It has been suggested that schizophrenia is related to the brain’s reduced capacity to integrate information across the different regions (Kahn et al., 2015; Stephan et al., 2006). The reduced capacity to integrate information has been associated with several phenomena in schizophrenia, including reductions in functional and structural connectivity, and reductions in grey and white matter volumes. The most reported deficit is lower global functional connectivity between many regions, including: subcortical regions; and the frontal, temporal, and occipital cortices. However, a replicated exception to this trend is increased functional connectivity between the thalamus and somatosensory and motor areas (Argyelan et al., 2014; Damaraju et al., 2014; Giraldo-Chica and Woodward, 2017; Skudlarski et al., 2010; Tu et al., 2015). Reduction in functional connectivity is suggested to be the result of alterations in brain structural connectivity at different levels from impaired synaptic plasticity (Friston, 1998) to reduction in the capacity of the structural connections at macroscale (Kahn et al., 2015; van den Heuvel et al., 2010). Reductions in grey and white matter volumes have been also reported across whole brain including thalamus, frontal, temporal, cingulate and insular cortex in patients with SZ (Ellison-Wright and Bullmore, 2010; Kahn et al., 2015; Segall et al., 2009; Staal et al., 2001). Our approach adds another piece to this global deficient phenomena and reveals for the first time that there is also a transient reduction in the activity patterns of FDs. Furthermore, reduced functional state connectivity within FMs is in agreement with the hypoconnectivity observed in previous studies among brain regions. While we did not investigate functional connectivity between brain regions, we observed decreased functional state connectivity between subcortical and somatosensory and somatomotor domains within the FMs, which could be an important window into a link between increased functional connectivity among these regions and decreased functional connectivity with the rest of the brain.

We propose that our approach is well-suited to examine the alterations in brain’s capacity to integrate information because it models the brain as a hierarchical functional architecture in which elements of each level of the hierarchy constructed from integrating the information of the lower level. This proposition was also supported by our findings. There is substantial evidence that there are distinct patterns for schizophrenia as detected by our analysis. In our analysis, the most affected regions and domains include the thalamus of the subcortical domain; BA 38 of the attention domain; the left insula, left BA 44, right BAs 41/42 and the rostral area of left BA 22 of the language domain; and the fusiform gyrus, medioventral, and lateral occipital cortex of the visual domain. The thalamus is known as a major brain structure affected both structurally and functionally in patients with SZ (Cheng et al., 2015; Damaraju et al., 2014; Giraldo-Chica and Woodward, 2017). Disruption in attention associated areas is frequently reported in patients with schizophrenia (Bowie and Harvey, 2006). Particularly, temporal pole area (BA38) is a key part of the theory of mind (ToM) network, which is classically impaired in patients with SZ and autism-spectrum disorder (Assaf et al., 2010). Furthermore, BAs 41/42 is primary auditory cortex, and together with the BA22 (auditory association cortex/Wernicke’s area), has been repeatedly implicated in the pathophysiology of auditory hallucinations in schizophrenia (Barta et al., 1990; Gavrilescu et al., 2010; Shinn et al., 2013; Vercammen et al., 2010). Alterations in the visual domain have been also observed as ocular convergence deficits (Bolding et al., 2012) and reduce amplitude of low-frequency fluctuations (ALFF) was observed across visual areas including the cuneus and lingual gyrus (Hoptman et al., 2010). Therefore, our findings are further buttressed by previous literature, suggesting spatial dynamics can provide a new dimension/level of schizophrenia-related alterations in the brain, which can potentially be leveraged to characterize clinical features in other patient groups.

## Conclusion

We proposed a novel framework that, for the first time, exploits the well-accepted brain functional hierarchical model to capture the spatial dynamics of brain functional organization. The present work reveals strong evidence that FDs evolve spatially over time including a broad spectrum of changes in regional associations from strong coupling to complete decoupling. Additionally, given that the brain reorganizes its activity at different interacting spatial and temporal scales, our hierarchical framework opens a new avenue to evaluate spatiotemporal variations within and between levels of the brain functional hierarchy providing a broader perspective of how the brain naturally functions. Preliminary assessments of the approach using healthy controls and patients with SZ demonstrates the ability of the approach to obtain new information of the brain function and detect alterations among patients with SZ. However, further investigations using different datasets and various cohorts should be performed to evaluate the benefits of studying spatiotemporal variations of brain functional domains for both basic and clinical neuroscience applications.

## Acknowledgments

This work was supported by grants from the National Institutes of Health grant numbers 2R01EB005846, R01REB020407, and P20GM103472; and National Science Foundation (NSF) grant 1539067 to Dr. V.D. Calhoun; and the National Institute of Mental Health grant number R01MH058262 and the Department of Veterans Affairs Senior Research Career Scientist award I01 CX0004971 to Dr. J.M. Ford.

